# An image segmentation pipeline optimized for human microglia uncovers sources of morphological diversity in Alzheimer’s disease

**DOI:** 10.1101/2024.02.01.577128

**Authors:** Robert M. De Jager, Annie J. Lee, Alina Sigalov, Mariko Taga

**Affiliations:** Center for Translational & Computational Neuroimmunology, Department of Neurology and the Taub Institute for Research on Alzheimer’s Disease and the Aging Brain, Columbia University Irving Medical Center, New York NY

## Abstract

Alzheimer’s disease (AD) is a neurodegenerative disorder characterized by cognitive decline and the accumulation of beta-amyloid plaques and neurofibrillary tangles. Microglia, the resident immune cells in the brain, play a crucial role in AD pathology, particularly in relation to amyloid plaques. However, the exact role of plaque-associated microglia and their morphological changes in AD progression remains under debate. In this study, we aimed to establish an automated image segmentation and analysis pipeline optimized for the detection of human microglia, and we assessed its utility by systematically investigating topological relationships between microglia morphology and amyloid plaques.

We accessed post-mortem brain tissue samples from persons with AD and utilized immunofluorescence staining to label microglia (IBA1) and amyloid plaques (Aβ1-42). The acquired images were processed using CellProfiler, an open-source image analysis software, to automatically segment microglia and measure their morphological features. Specifically, since activated (stage III) microglia have a condensed morphology with retracted processes, we prioritized a morphological parameter called “compactness” in our analyses to capture shape changes in microglial found in proximity to amyloid plaques.

Our results revealed that microglia unassociated with plaques (Mi^p-^) were more abundant than microglia associated with plaques (Mi^p+^) in the Dorsolateral Prefrontal Cortex (DLPFC) of both men and women with AD. Furthermore, we observed that Mi^p+^ exhibited a significantly more ramified shape and had a higher expression of the IBA1 microglial marker gene compared to Mi^p^. There were no significant differences in microglia morphology between men and women.

Our study highlights the utility of automated image analysis in characterizing detailed microglia morphology at the single-cell level and its relationship with AD pathology.

## Introduction

Alzheimer’s disease (AD) is the most common form of dementia in which cognitive function gradually declines with advancing age. The pathology of AD is characterized by the presence of two types of protein aggregates: beta-amyloid plaques^1^ (Aβ_1-42_) and neurofibrillary tangles ^2^. Several cell types are involved in the formation of amyloid plaques. Many studies focus on microglia, the resident immune cells in the central nervous system, due to their close association with Aβ_1-42_ plaques. However, the role of plaque-associated microglia in the progression of AD remains incompletely understood, especially in humans ^3–5^. Microglia are reported to be involved in the degradation and internalization of beta-amyloid plaques ^6,7^, and impaired or altered microglial responses to beta-amyloid plaques are associated with an increased risk of AD ^8,9^. There is evidence to suggest that microglia acquire different morphologies and activation states in AD^10^. Under homeostatic conditions, microglia have fine and highly ramified processes, and they are described as “surveilling” the brain parenchyma. In various pathological conditions, microglia become activated which is manifested by an amoeboid cell form^10,11^ with retracted processes and a swollen cell soma^10,12^. Prior studies analyzed data collected by human operators who assigned each microglia one of three basic shapes based on observing the tissue under the microscope: stage I cells are larger with extensive processes, stage II cells are round with plum cytoplasm with thicker processes and stage III cells are rounder with thick, retracted processes. Interestingly, higher numbers of stage III microglia were associated with cognitive decline and AD pathologies^10^. These findings suggest that microglia morphology seems to be related to an important role in AD progression, although its assessment remains rudimentary. In most post-mortem human brain studies, morphology is often analyzed manually, which limits the number of microglia that can be analyzed. Furthermore, the choice of the measurement of microglial shape from the number and thickness of processes and soma shape can be biased. Therefore, it is necessary to develop a high-throughput automated image analysis tool that characterizes microglia morphology with the exact measurement of cellular shape to expand the possibility of understanding the relationship between different microglia states and AD pathology. This study investigates the spatial association between microglial phenotype and one of the main forms of AD pathology, amyloid plaques, using the open-source analysis software CellProfiler^13^.

Cellprofiler is an image analysis software designed to extract quantitative measurements from cellular structures, and the software provides 17 shape measurements which include the area, the perimeter, and the compactness. The aim of this study was to investigate the spatial relationship between microglia morphology and amyloid plaques, by developing an automated image analysis pipeline capable of automatically segmenting microglia and then extracting morphology measurements from immunofluorescence images generated from human post-mortem fixed tissues. We hypothesized that a measure known as “compactness”, calculated as the mean squared distance of an object’s pixel from its centroid divided by the area of the object, would accurately capture the changes in the microglial shape associated with amyloid plaques. This study’s findings could have a significant impact on scaling up investigations of microglial morphology related to AD pathology and other diseases by utilizing our automated image analysis pipeline. The current approach of relying on human operators to manually score images is known to be slow, inefficient, and susceptible to variations in interpretation among different operators. Therefore, the deployment of automated image segmentation and analysis software will enhance the efficiency and reliability of our experimental pipeline. As a result of this approach, we may be able to better characterize the changes in the microglial shape associated with the disease and to develop new targets to reverse the changes in microglial shape and slow down the progression of the disease.

## Material and Methods

### Post Mortem Human brain Tissues

We accessed formalin-fixed post-mortem brain tissue samples from RUSH University Medical Center and New York Brain Bank (NYBB) who fulfilled the criteria for a diagnosis of AD using the Reagan criteria^14^. All participants consent to brain donation at death. The demographic characteristics of these individuals are summarized in **Supplementary Table 1**.

#### ROSMAP cohort

The Religious Orders Study (ROS) and Memory and Aging Project (MAP) are both cohort studies directed by Dr. David Bennett at RUSH University. Individuals over the age of 65 were free of dementia at enrollment and consented to brain donation at the time of death.

Both cohorts conduct the same annual assessments, which include 19 cognitive functional tests, using validated procedures to diagnose AD and other dementias. All participants undergo a standardized structured assessment for AD which includes CERAD, Braak Stage, NIA-Reagan, and a global measure of AD pathologic burden with modified Bielschowsky. Additionally, amyloid load and PHF Tau tangles are measured using immunocytochemistry, image analysis, and stereology. Both frozen and fixed brain tissues are accessible upon request from these subjects.

#### New York Brain Bank

The New York Brain Bank (NYBB) at Columbia University banked brain tissues coming from the Columbia ADRC (Alzheimer’s Disease Research Center), the NIA Alzheimer’s disease Family Based Study (NIA-AD FBS), the Washington Heights, Inwood Columbia Aging Project (WHICAP), and the Biggs Insitute Brain Bank at the University of Texas Health Science Center at San Antonio. The Columbia ADRC cohort consists of clinical participants in the Columbia ADRC who have consent for brain donation. For all ADRC brain donations, one hemisphere is fixed while the other one is banked fresh and stored at -80°C. The NIA-AD FBS has recruited and followed 1,756 families from various ethnic groups with suspected late-onset AD, which include 9,682 family members and 1,096 unrelated, non-demented elderly. This initiative provides valuable biological resources such as brain tissue, plasma, and peripheral Blood Mononuclear Cells (PBMCs) along with genetic data. The Resource Related Cooperative Agreement extended its recruitment efforts to include familial early-onset AD. Since 1992, the WHICAP has enrolled more than 6,000 participants, with representative proportions of African Americans (28%), Caribbean Hispanics (48%), and non-Hispanic whites (24%). Finally, the Biggs Institute Brain Bank at the University of Texas Health Science Center collects postmortem brain, spinal cord, cerebrospinal fluid, and dermal tissue from the study participants. The consent for the brain tissues was obtained from the donor’s legal next-of-kin before the autopsy. The left half hemisphere is fixed in 10% neutral-buffered formalin and the right hemisphere is banked fresh at -80°C. After a minimum of 4-week fixation period, the fixed tissue is sectioned and sampled in accordance with National Institute on Aging-Alzheimer’s Association AD neuropathologic guidelines.

### Immunohistochemistry

Six μm sections of formalin-fixed paraffin-embedded (FFPE) tissue from the dorsolateral prefrontal cortex (Brodmann Area 9) were de-paraffinized with CitriSolv (Xylene substitute, VWR) for 20 min. Heated-induced epitope retrieval was performed with citrate (pH=6) using a microwave (800W, 30% power setting) for 25 min. The sections were blocked with blocking medium (3% BSA) for 30 min at Room Temperature, then incubated with primary antibody anti-IBA1 (Wako; 011-27991; dilution 1:100) and anti-Amyloid Beta_1-42_ (Biolegend; 805501; 1:500) prepared in 1% BSA for overnight at 4°C. Tissue Sections were washed three times with PBS and incubated with fluorochrome conjugated secondary antibodies (Thermo Fisher, dilution 1:500) for one hour at RT. After three times washing, the sections were exposed to True Black Lipofuscin Autofluorescence Quencher to minimize the autofluorescence. Anti-fading reagent with Dapi (P36931, Life technology) was used for coverslipping. For each subject, 40 images of cortical grey matter at magnification x20 (Nikon Eclipse Ni-E fluorescence microscope) were taken in a zigzag sequence along the cortical ribbon to ensure that all cortical layers are represented in the quantification in an unbiased manner. We processed the acquired images using CellProfiler v4.2.1 for automated segmentation.

### Automated microglia segmentation using the CellProfiler software

Using CellProfiler v4.2.1, we developed an extensive pipeline for segmenting microglia and measuring their morphology. Using the “IdentifyPrimaryObjects” module, we initially identified and segmented nuclei (DAPI). Next, the “EnhanceOrSuppressFeatures” module was employed to enhance microglia branches to facilitate the detection of microglia morphology. Subsequently, the IBA1 image underwent using the “IdentifyPrimaryObjects” module. To consolidate small microglia branches, the “SplitOrMergeObjects” module was utilized, with a maximum distance of 1 pixel specified for merging objects. This module enables the merging of two or more unconnected components. In order to specifically detect “real cells”, segmented IBA1 objects were related to segmented nuclei (DAPI) to filter IBA1+DAPI+ using the “RelateObjects” module. Therefore, only microglia positive for nuclei will be used for the analysis. The shape and intensity of IBA1+DAPI+ were measured using “MeasureObjectSizeShape” and “MeasureObjectIntensity” modules.

Amyloid plaques were segmented using the “IdentifyPrimaryObjects” module, and small split objects were combined into a single object using the “SplitOrMergeObjects” module. The “RelateObjects” module was used to relate IBA1+ DAPI+ cells with segmented amyloid plaques. IBA1+ DAPI+ cells associated with amyloid plaques were classified as microglia associated with plaques, and IBA1+ DAPI+ cells unrelated to amyloid plaques were classified as microglia not associated with plaques.

The morphological parameters of each joint-segmentation-defined cell were measured with the MeasureObjectSizeShape module and all data was exported with the ExportToSpreadsheet module. A total of 29 morphological features were extracted for each IBA+DAPI+ cell. According to prior work^5^, the “Object_compactness” morphology parameter was prioritized. 11,399 segmented microglia (IBA1+DAPI+ cells) have been collected using CellProfiler software from NYBB and ROSMAP cohorts.

### Statistical analyses

We conducted our statistical analyses primarily using logistic regression for each sample obtained from NYBB and ROSMAP. Subsequently, a meta-analysis of the results from both ROSMAP and NYBB was carried out using a fixed-effects model with the inverse-variance weighted average method using the R package metafor. All statistical analyses were performed using R 4.1.3 (https://www.r-project.org/).

## Results

### Study design

We accessed AD brain tissue from two different sample collections to evaluate the robustness of our approach: (1) The Religious Order Studand Memory & Aging Project (ROSMAP) resource consisting of prospectively collected brains from participants in two cohort studies of cognitive aging and (2) the New York Brain Bank, where we accessed samples in their AD collection. Demographic information is presented in **Supplementary Table 1**; all participants fulfill a pathologic diagnosis of AD by Reagan criteria. 60% of the participants are female in ROSMAP and 62.5% in NYBB. From each collection, we obtained a 6 μm section and stained it with anti-IBA1 and Amyloid Beta_1-42_. Images were captured on a Nikon Eclipse Ni-E fluorescence microscope as outlined in the Methods section at 20x resolution, and images were segmented using a customized pipeline deployed in CellProfiler version v4.2.1 (see Methods) (**Figure 1A and B**). The resulting microglia were then classified as microglia associated with plaques (Mi^p+^) if at least one pixel of the microglia was juxtaposed to one pixel of an amyloid plaque; all other microglia were classified as microglia unassociated with plaques (Mi^p-^).

**Figure 1:**
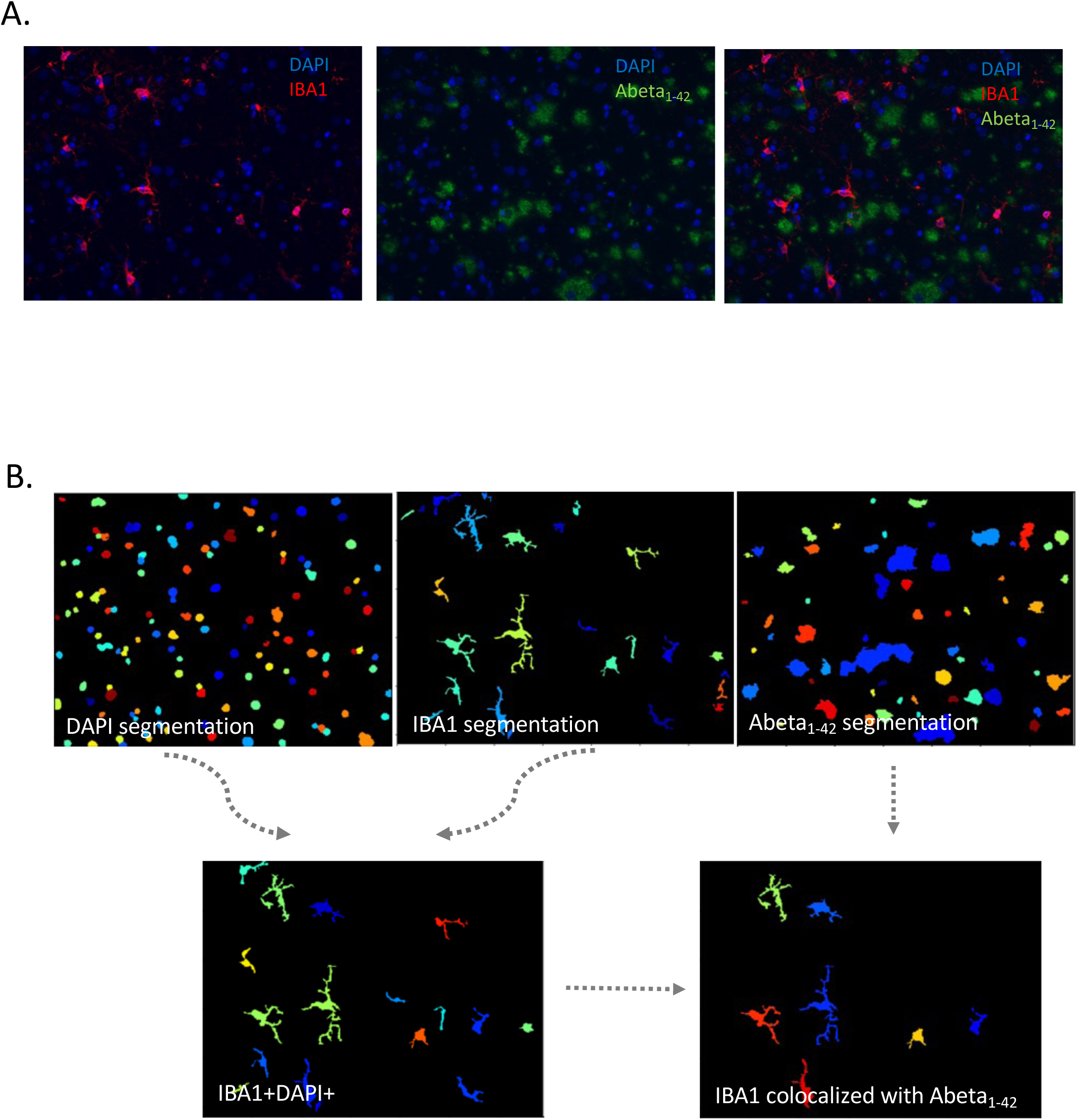
**(A)** Immunohistochemistry was performed on human post-mortem brain tissues using IBA1, a general marker of microglia (in red), and Abeta_1-42_ to detect amyloid plaques (in green). Images were acquired at a magnification of 20X using an immunofluorescence microscope. **(B)** Microglia and Abeta_1-42_ plaques were segmented using a segmentation pipeline created with CellProfiler Software.

### The neocortex of persons with AD contains more microglia that are unassociated with plaques in both women and men

We first compared the frequency of Mi^p+^ and Mi^p-^ cells in each sample obtained from the two different sample collections by fitting a logistic regression for each sample. The result showed a negative beta (b) in each of the two sample collections (ROSMAP and NYBB), indicating that there are significantly fewer Mi^p+^ compared to Mi^p-^ in persons with AD (**Figure 2A**). A meta-analysis of results from ROSMAP and NYBB was conducted using a fixed-effects model with an inverse-variance weighted average and returned a highly significant result (p<0.0001).

**Figure 2:**
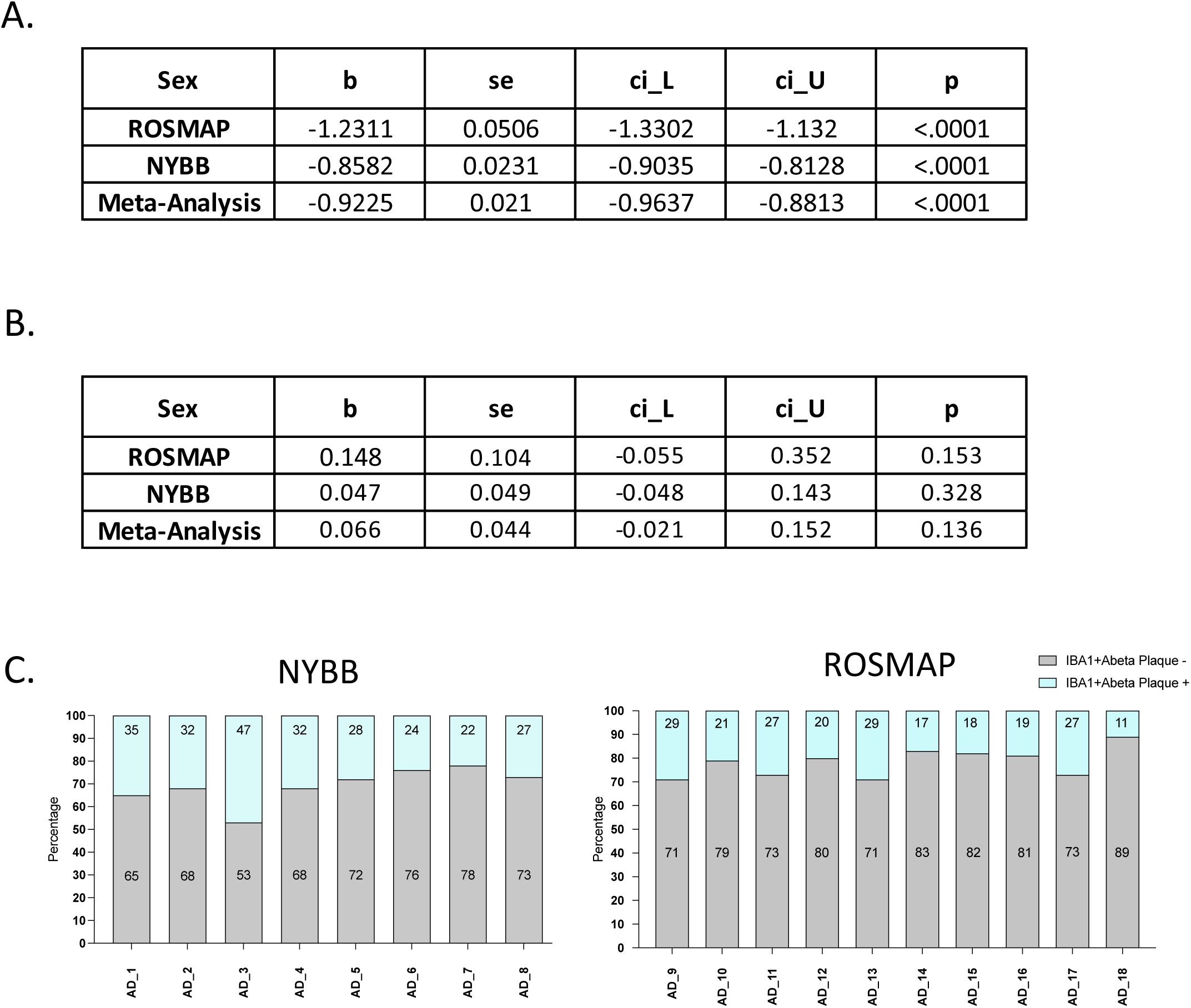
**(A)** Comparison of Mip- and Mip+ frequencies in AD subjects across two cohorts. The quantification of the number of Mip- and Mip+ was performed using the CellProfiler pipeline, showing comparable results between ROSMAP (n=10) and NYBB (n=8). The threshold of significance is p<0.05. **(B)** Comparison of the frequency of Mip^-^ and Mip^+^ between women and men. Mip+ and Mip-frequencies are similar between both genders and comparable results are reported between ROSMAP (n=10) and NYBB (n=8). The threshold of significance is p<0.05. **(C)** Percentage of Mip^+^ and Mip^-^ in AD subjects from ROSMAP and NYBB.

Since women are more likely to develop AD than men^15^, we evaluated the effect of sex on the frequency of Mip^-^ and Mip^+^ cells by fitting a logistic regression model (**Figure 2B**). Our approach consisted of performing this analysis separately for ROSMAP (p=0.153) and NYBB samples (p=0.328), and then we conducted a meta-analysis of the results from ROSMAP and NYBB samples using inverse-variance weighted averages with fixed-effects models. The result showed that there was no significant difference in the frequency of the Mip^-^ and Mip^+^ cells between men and women (p=0.136).

### Microglia associated with amyloid plaques have a more “ramified” shape than microglia non-associated with amyloid plaques

We tested our hypothesis that our automated image analysis pipeline yields quantitative measures of microglia morphology that will enable the study of microglial shape in relationship to AD. Because a minority of rounded microglia have been previously found to be associated with AD^10^, we chose the “compactness” measure provided by CellProfiler as our primary outcome to in assessing the change in microglial shape associated with amyloid plaques. “Compactness” is defined as the mean squared distance of an object’s pixels from the centroid of the object, divided by its area. A circular object has a compactness of 1, and irregularly shaped objects or objects with holes (such as ramified microglial cells) having a compactness >1.

To visualize the distribution of compactness measures among the microglia of each AD individual from both cohorts, we plotted histograms that indicate that, consistently, the distribution of compactness scores for the microglia derived from each tissue section has a peak of cells with a relatively low compactness score (meaning that they are rounder) and a long tail of cells with more complex shapes (**Figure 3 A and B**).

**Figure 3:**
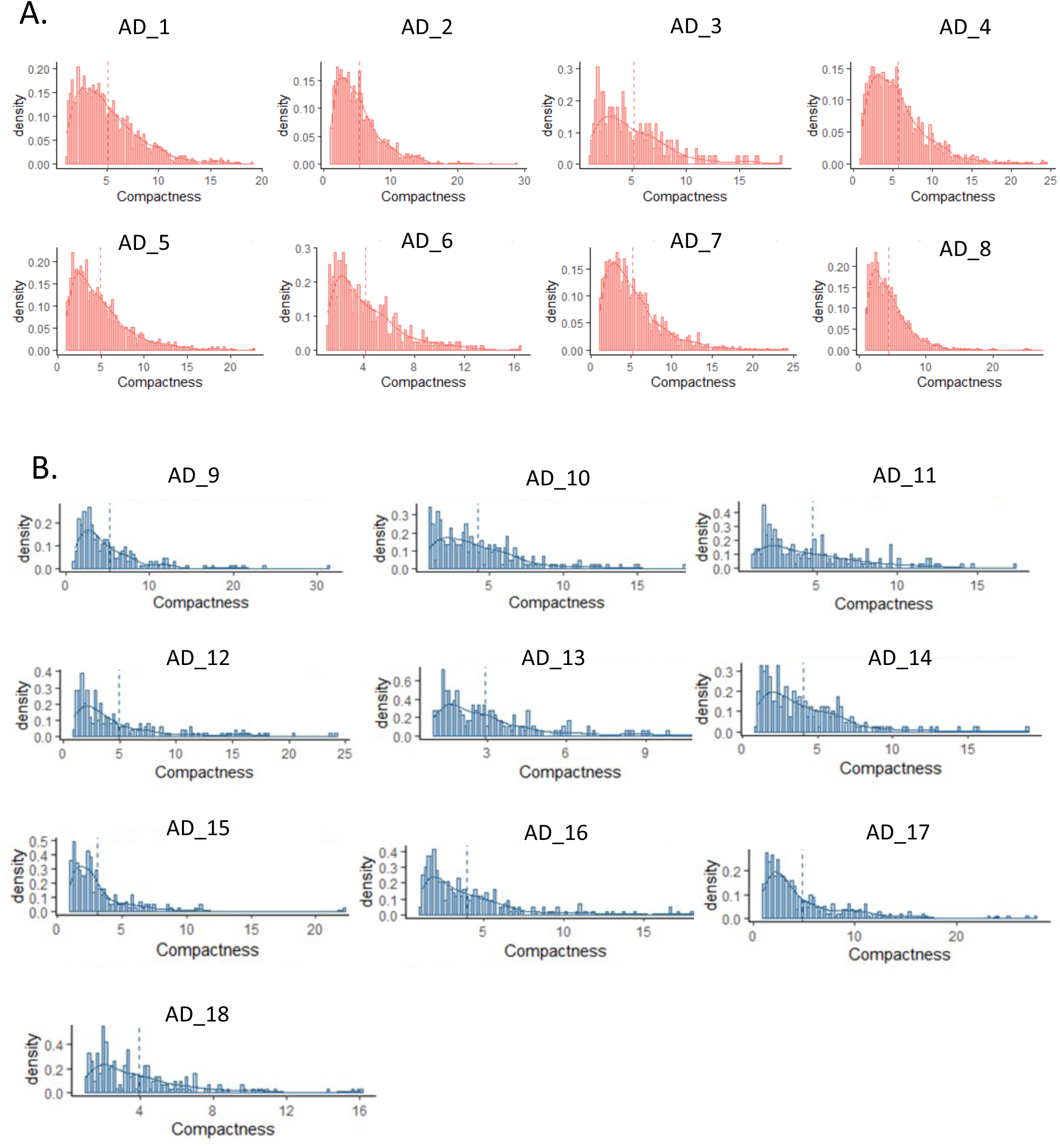
Distribution of compactness measures obtained from CellProfiler among all microglia (Mip^+^ and Mip^-^) of each AD individual from **(A)** NYBB (n=8) and **(B)** ROSMAP (n=10) using a histogram.

To compare the compactness of Mi^p+^ and Mi^p-^ cells, we fitted a logistic regression for each sample; thus, we have analysis and results for each tissue section/participant. We then conducted a meta-analysis combining results for all participants within one of the sample collections using a fixed-effects model with an inverse-variance weighted average method. We thus performed separate meta-analyses steps for ROSMAP and NYBB, and we finally conducted a second meta-analysis to combine ROSMAP and NYBB results through a fixed-effects model with an inverse-variance weighted average method (**Figure 4 A and B**). The result showed that, in each tissue section from an individual participant, Mi^p+^ cells have a has a higher value of compactness compared to Mi^p-^ cells (positive effect of plaque on compactness). This observation is significant in both sample collection and in the final analysis combining all data (p<0.001). Thus, Mi^p+^ are less rounded and have more ramified shapes than those microglia that are not in contact with amyloid plaques.

**Figure 4:**
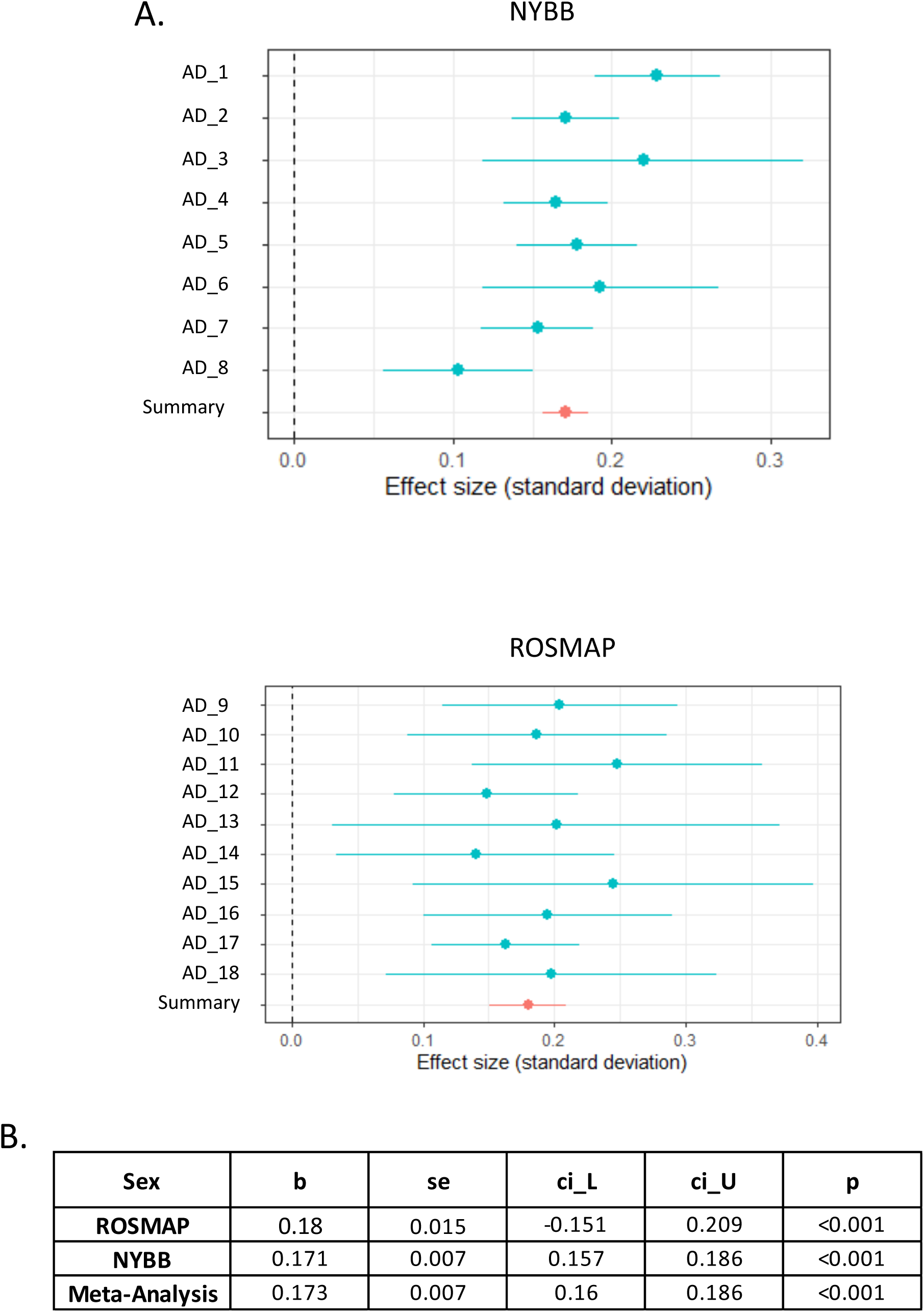
**(A)** Forest plots illustrating the association of compactness between Mip+ vs Mip- in each AD individual from NYBB (n=8) and ROSMAP (n=10). The effect of Mi^p+^ on the compactness is comparable between both cohorts (positive effect). **(B)** Meta-analysis across NYBB and ROSMAP samples with a significance threshold of p<0.05.

### No sex-specific differences in compactness between Mi^p+^ and Mi^p-^ cells

In order to investigate whether microglia compactness differs between men and women, we repeated our analyses by meta-analyzing the data for men and women separately in each sample collection. The difference in the association between men and women was then tested by fitting a single fixed-effects meta-regression model with sex as a moderator. We then summarized results by performing a meta-analysis of results from both ROSMAP and NYBB through a fixed-effects model with the inverse-variance weighted average method (**Figure 5 A and B**). The result showed that the compactness value is significantly higher in Mi^p+^ cells than Mi^p-^ cells in both women and men (positive effect of plaque on compactness); and while men appear to have, on average, greater compactness in Mi^p+^ cells than women(**Figure 5 A and B**), this difference is not statistically significant (p=0.097). We note that this trend may well become significant in larger sample sizes since it is observed in both sample collections; we, therefore, interpret our results as suggesting that there are no sex-based differences in microglial compactness relative to amyloid plaques of large effect in human AD samples. However, an effect of modest size cannot be excluded at this time.

**Figure 5:**
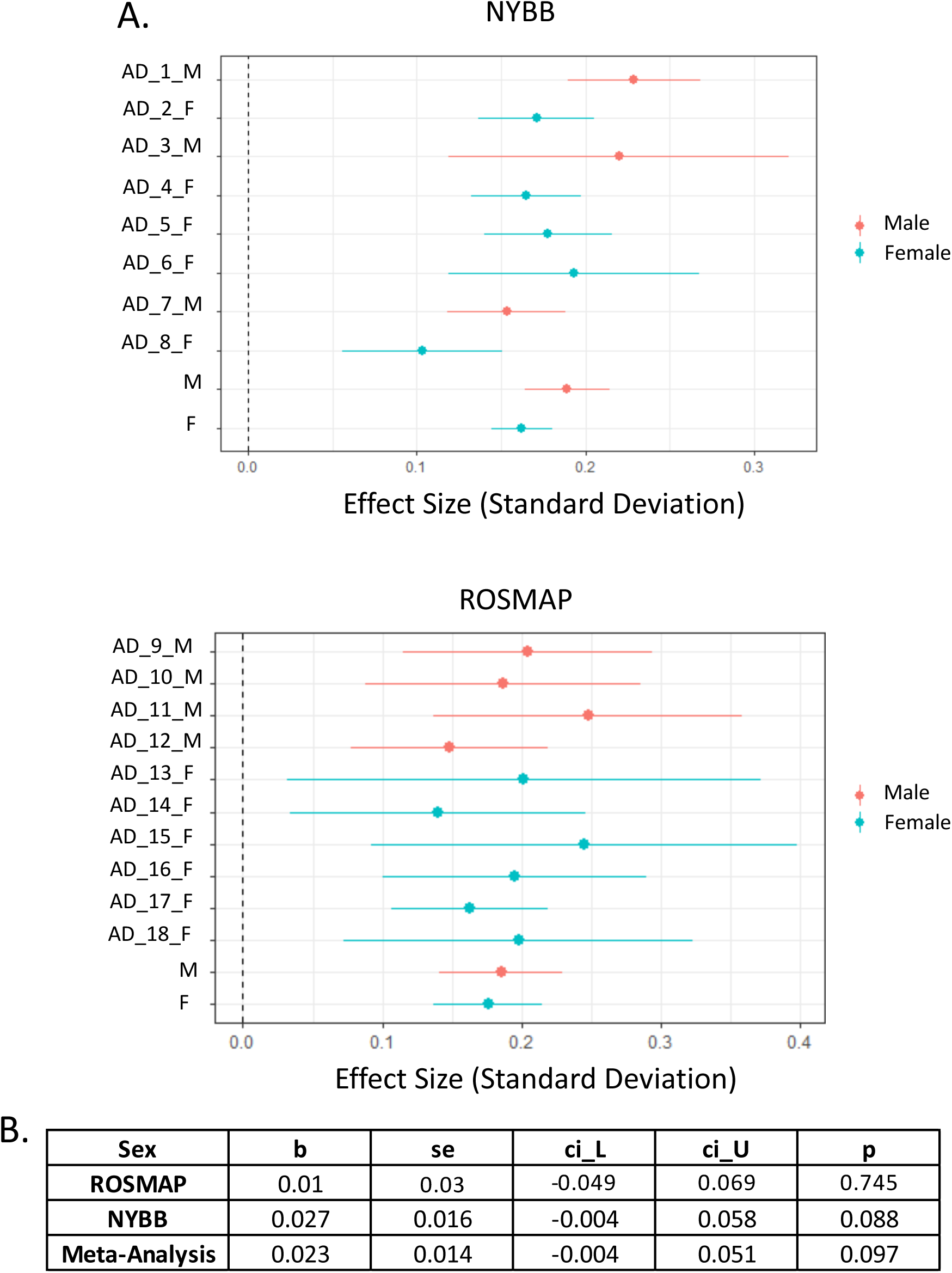
**(A)** Forest plots illustrating the effect of sex on Mip^-^ and Mip^+^ compactness within each AD individual from NYBB (n=8) and ROSMAP (n=10). The effect of sex on Mip- and Mip+ compactness is comparable between both cohorts (positive effect). **(B)** Meta-analysis across NYBB and ROSMAP samples with a significant effect of sex on Mip^+^ and Mip^-^ compactness.

### IBA1 expression is increased in ramified Mi^p+^ cells

Finally, we investigated the potential variation in IBA1 expression based on the microglia morphology. In this study, we compared the association of compactness to the intensity of IBA1 by fitting a linear regression for each sample. The compactness score was transformed to approximate a normal distribution. A meta-analysis was conducted across the association studies for Mi^p+^ and Mi^p-^ cells separately. The difference in the association between Mi^p+^ and Mi^p-^ was tested by fitting a single fixed-effects meta-regression model with plaque status as a moderator.

These analysis steps were performed for each ROSMAP and NYBB sample separately and then meta-analyzed in each sample collection separately. Finally, results from the two separate sample collections were meta-analyzed to produce the final results. A fixed-effects model with inverse-variance weighted average method was used in these stepwise analysis approach. **Figure 6 A** indicates that for both ROSMAP and NYBB samples, there are positive correlations between compactness and intensity of IBA1 in both Mi^p+^ and Mi^p-^ cells (p=1.56×10^-9^ and p=1.93×10^-2^ respectively). Interestingly, after meta-analysis, we observed that the effect of the intensity of IBA1 on compactness was significantly higher in Mi^p+^ cells compared to Mi^p-^ cells (**Figure 6 B**) (p=6.48×10^-8^). These findings suggest that microglia associated with plaques have higher IBA1 expression than microglia non-associated with plaques.

**Figure 6:**
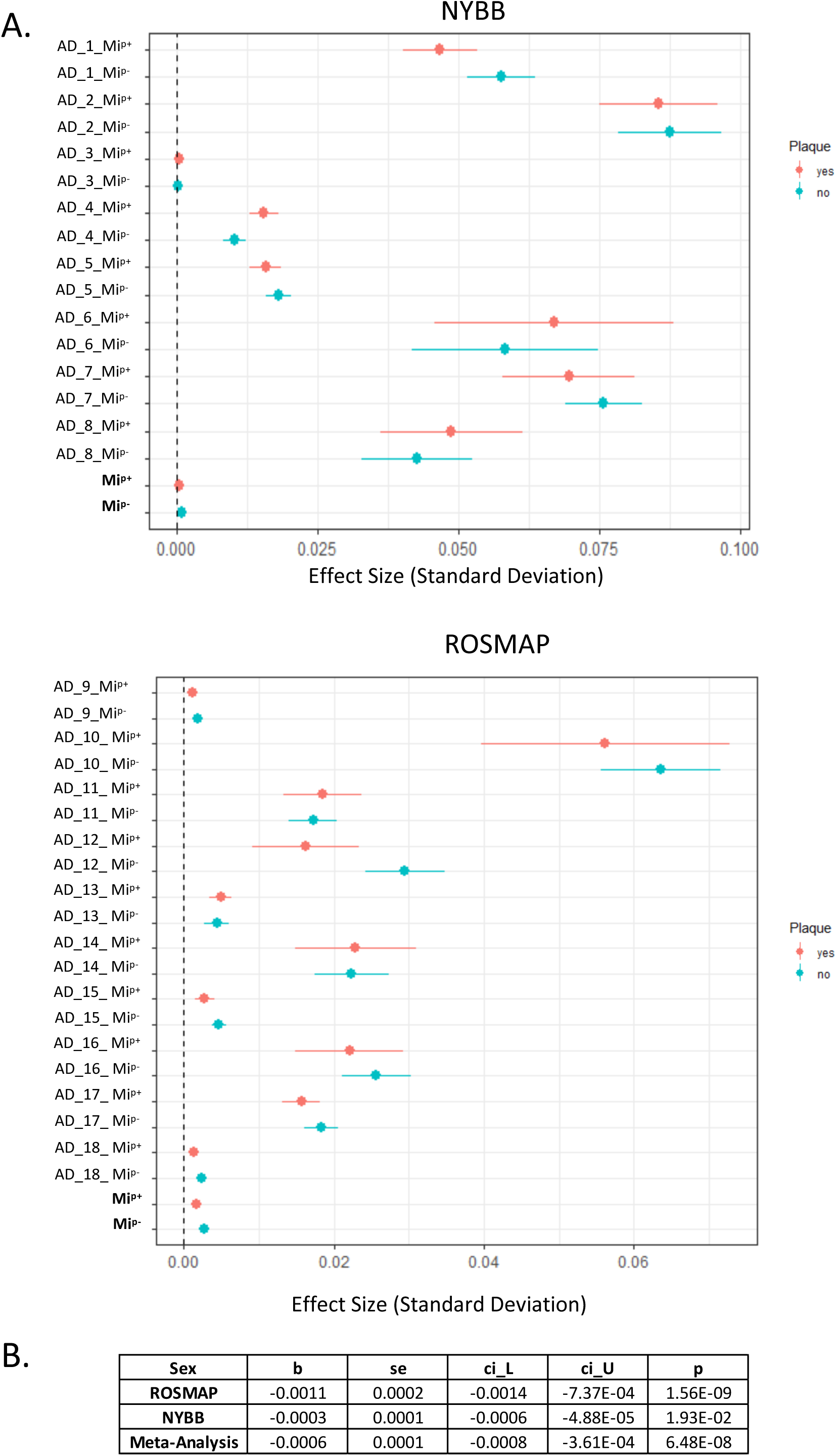
**(A)** Forest plots illustrating the association of Mip^-^ and Mip^+^ compactness to IBA1 intensity in each AD individual from NYBB (n=8) and ROSMAP (n=10). The effect of Mip^-^ and Mip^+^ compactness on IBA1 intensity is comparable between both cohorts (positive effect). **(B)** Meta-analysis across NYBB and ROSMAP samples with a significant threshold of p<0.05.

## Discussion

In this study, we have established a robust and comprehensive image segmentation and downstream analysis pipeline capable of being applied to two distinct brain sample collections. Results are consistent in the two collections, demonstrating the robustness of our image segmentation pipeline despite many differences in tissue acquisition, preparation, fixation, and storage between the two sample collections. Technical differences in tissue quality and properties (such as background autofluorescence) are a major challenge in deploying automated image segmentation pipelines across different sample collections. Here, our primary objective was to (1) develop an automated method to accurately capture and quantify microglial morphology that avoids the limitations of manual scoring by human operators and (2) capture protein expression patterns as well as topological characteristics relative to important landmarks such as amyloid plaques. Our approach presents a versatile and adaptable methodology that can be effectively applied to address a wide range of inquiries pertaining to human microglia. Here, we successfully demonstrated that capturing data at the single-cell level and retaining topological information allows us to powerfully interrogate hypotheses, even when working with a relatively small number of samples. Our study included 18 participants with a pathologic diagnosis of AD, and we acquired data on 11,399 microglia, resulting a substantial statistical power despite small sample sizes. Our analyses revealed substantial morphological heterogeneity among microglia in

AD brains. The preservation of topological information was a key component in resolving two subsets of microglia defined by their physical relationship to amyloid plaques. Notably, microglia associated with amyloid plaques exhibited higher compactness scores, indicating a more complex shape with projections away from the main cell mass. Interestingly, this finding suggests that the rounded cells with condensed cytoplasm and projections that have been noted to increase in frequency with AD are located away from the amyloid plaques. Our observation is in agreement with studies that examined the activation and morphology of microglia in association with amyloid plaques using the immunohistochemistry approaches. However, it is important to note that the majority of these findings relied on staining observations rather than precise morphology measurements ^16–18^. Ramified microglia have been frequently observed in diffuse plaques, which were found to be the predominant type of plaques among the amyloid plaques^16^. Prior studies have also reported a spatial association between plaques devoid of tau-positive structures and ramified microglia^17,18^, suggesting that amyloid plaques don’t trigger microglial activation. Furthermore, our observations align with our prior modeling that stage III microglia might be involved in the accumulation of Tau protein rather than amyloid proteinopathy ^10^. Traditionally, ramified microglia were qualified as being in a “resting” state. However, a study employing two-photon microscopy in mice has redefined these cells as being in a “surveillance” state characterized by their dynamic motility^19^. This is consistent with the observation of increased IBA1 expression with a greater compactness score (more ramified shape), as this protein is involved in cell skeleton reorganization, microglia motility^20^, and phagocytosis^21^. The presence of this notable interaction between microglial types (Mi^p+^ and Mi^p-^) and the association of IBA1 intensity with increasing compactness value supports the concept that Mi^p+^ and Mi^p-^ cells represent distinct microglial subpopulations with functional differences. Now, the current challenge is to align the topologically defined subtypes of microglia, which exhibit morphological and protein expression variations, with the subtypes identified through single-cell transcriptomics such as disease-associated microglia that have been associated with AD pathologic features ^22–24^.

Our study had several limitations, with the most significant one being its limited sample size. However, we implemented a unique study design that involved collecting extensive data at the single-cell level within each individual in two different sample collections. This approach enabled us to conduct intra-individual analyses followed by a meta-analysis across individuals and sample collections, thereby enhancing our statistical power compared to averaging measures across all cells within each individual. Such a per-individual summary score has been the traditional approach for manually segmented studies and other approaches for image segmentation that obtain a summary measure of immunofluorescence for each image. Our approach provides much more granular and scalable data that set the stage for well-powered investigations of microglial morphology and topology in human tissue.

Overall, segmentation of morphologically complex and diverse cells such as microglia remains a major challenge, particularly as it must be conducted in the topologically complex cytoarchitecure of the neocortex, which has 6 well-defined layers and a host of different micro-environments around vascular and meningeal compartments as well as a variety of pathological features such as aging-related proteinopathies. Further, application of rigorous statistical methodology is essential to produce robust, reproducible results, as illustrated here, and such methodologies and reduced variance in morphological measures from systematic assessments wil help to minimize sample sizes in future studies. With our image pre-processing and analytic pipelines in place, we can now attack more complex questions relating to microglial pathology, and the scability of our approach enables the development of challenging study designs such as genetic studies that typically require large sample sizes.

---

**Supplementary Table 1:** Demographic characteristics of the cortical tissue samples (n=18) obtained from two brain collections: New York Brain Bank (NYBB) and ROS/MAP (ROS/MAP: n=10, NYBB: n=8) for the immunohistochemistry staining. The AD pathology is defined using NIA-Reagan neuropathologic criteria.

|  | Gender | Age at death | Pathological Diagnosis | Cohort/Brain bank |
| --- | --- | --- | --- | --- |
| MP10-60 | M | 69 | AD | NYBB |
| MP11-189 | F | 89 | AD | NYBB |
| MP13-9 | M | 79 | AD | NYBB |
| MP13-101 | F | 89 | AD | NYBB |
| MP14-225 | F | 89+ | AD | NYBB |
| MP15-239 | F | 68 | AD | NYBB |
| MP17-91 | M | 71 | AD | NYBB |
| MP17-114 | F | 88 | AD | NYBB |
| 10490993 | M | 86 | AD | ROSMAP |
| 10518782 | M | 87 | AD | ROSMAP |
| 11302830 | M | 85 | AD | ROSMAP |
| 15113169 | M | 84 | AD | ROSMAP |
| 20665307 | F | 95 | AD | ROSMAP |
| 21181988 | F | 92 | AD | ROSMAP |
| 29933130 | F | 94 | AD | ROSMAP |
| 50401002 | F | 86 | AD | ROSMAP |
| 50403446 | F | 89 | AD | ROSMAP |
| 50410283 | F | 86 | AD | ROSMAP |

